# The DNA Methyltransferase DMAP1 is Required for Tissue Maintenance and Planarian Regeneration

**DOI:** 10.1101/2024.04.10.588909

**Authors:** Salvador Rojas, Paul G. Barghouth, Peter Karabinis, Néstor J. Oviedo

## Abstract

The precise regulation of transcription is required for embryonic development, adult tissue turnover, and regeneration. Epigenetic modifications play a crucial role in orchestrating and regulating the transcription of genes. These modifications are important in the transition of pluripotent stem cells and their progeny. Methylation, a key epigenetic modification, influences gene expression through changes in histone tails and direct DNA methylation. Work in different organisms has shown that the DNA methyltransferase-1-associated protein (DMAP1) may associate with other molecules to repress transcription through DNA methylation. Thus, DMAP1 is a versatile protein implicated in a myriad of events, including pluripotency maintenance, DNA damage repair, and tumor suppression. While DMAP1 has been extensively studied *in vitro*, its complex regulation in the context of the adult organism remains unclear. To gain insights into the possible roles of DMAP1 at the organismal level, we used planarian flatworms that possess remarkable regenerative capabilities driven by pluripotent stem cells called neoblast. Our findings demonstrate the evolutionary conservation of DMAP1 in the planarian *Schmidtea mediterranea*. Functional disruption of DMAP1 through RNA interference revealed its critical role in tissue maintenance, neoblast differentiation, and regeneration in *S. mediterranea*. Moreover, our analysis unveiled a novel function for DMAP1 in regulating cell death in response to DNA damage and influencing the expression of axial polarity markers. Our findings provide a simplified paradigm for studying DMAP1’s epigenetic regulation in adult tissues.

**Highlights:**

- Epigenetic regulation through DMAP1 is evolutionarily conserved in *Schmidtea mediterranea* and is crucial for tissue maintenance and regeneration.
- Neoblast differentiation into epithelial, muscle, digestive, and neural fate requires DMAP1.
- DMAP1 regulates DNA stability and cell death during adult cell turnover.
- DMAP1 regulates the spatial expression of axial polarity markers in *S. mediterranea*.

## Introduction

Embryonic development and events such as cellular turnover and regeneration at adult stages rely on a fine regulation of DNA transcription to ensure the correct transition of pluripotent stem cells and their progeny. Epigenetics, or the genetic changes that can alter dynamics of DNA transcription, and translation without changing DNA sequence, ensures the proper transition and differentiation of cells^1–3^. Specifically, methylation is one of the genetic modifications that can alter proper gene expression. One form by which methylation regulates proper gene transcription is through post-translational modifications acting on the histone tails and by methylating the DNA ^1,3^. Methyl groups are added to cytosine bases with a guanine next to them (CpG islands). DNA methyltransferase (DNMT) is a chief enzyme that maintains mammalian DNA methylation by having its N-terminus bind to DNA methyltransferase-associated protein 1 (DMAP1) and can establish a repressive transcription complex with histone deacetylases (HDACs)^2–11^.

DMAP1 does not only form a complex with DNMT1 but may associate with a variety of different complexes. For example, DMAP1 is also a member of the TIP60-p400 complex that maintains embryonic stem cell pluripotency^12–18^. Furthermore, DMAP1 is also necessary for enabling DNA damage repair^2,11,14,17,19–23^. The knockdown of DMAP1 function impairs DNA repair by altering the activation of ATM (Ataxia Telangiectasia Mutated), which is an essential regulator of the cellular response to DNA damage^21^. DMAP1 also plays a role as a mediator for tumor suppressors^9,11,15,24,25^ to induce the p53/p21 signaling cascade^21^. Given the complex epigenetic regulation of DMAP1, most studies on its function have been performed *in-vitro*.

To address the regulation of DMAP1 in the complexity of the adult body, we propose using planarian flatworms. Planarians are multicellular animals with a remarkable capacity to repair and replace lost structures through regeneration^26,27^. Planarians’ regenerative capacity relies on pluripotent stem cells called neoblast that are responsible for the maintenance of homeostasis^26,27^. Neoblasts are the only cells capable of proliferation and, therefore, responsible for propagating cellular progeny that differentiate and sustain the renewal and repair of many adult tissues. Thus, planarians provide an anatomically simple platform to analyze epigenetic regulation of cell function at the organismal level. Furthermore, planarians offer unprecedented opportunities to learn about epigenetic-mediated transcriptional regulation in invertebrates^28,29^. Recent work using planarians uncovered evolutionary conservation of epigenetic regulation and identified novel roles in tumor suppression, the neoblast transcriptional programs, adult stem cell pluripotency, and regeneration^25,29–34^. Previous research has also shown that the planaria *Schmidtea mediterranea* genome does not contain any cognate DNMTs, and the genome lacks cytosine-dependent methylation^29^.

Here we use *S. mediterranea* to analyze epigenetic regulation in planaria. Although previous studies suggested that *S. mediterranea* does not possess any DNMTs, we did, however, find an ortholog of DMAP1. We found that *Smed-dmap1* (will be referred to as *Dmap1* from here on) is evolutionarily conserved in planarians. Functional disruption of *Dmap1* with RNAi revealed critical roles for tissue maintenance, neoblast differentiation, and regeneration. Our analysis also identified that *Dmap1* regulates cell death in the presence of DNA damage and uniquely establishes a novel function for *Dmap1* as a regulator of expression of axial polarity markers in planaria. Altogether, our results present a simplified paradigm to analyze DMAP1 epigenetic regulation in adult tissues.

## Material and Methods

### Planarian Maintenance

All experiments were conducted using the planarian *Schmidtea mediterranea* asexual strain CIW4. Maintenance of animals was conducted as previously described^35^. Animals were starved for a week prior to use in experiments.

### Quantitative RT-PCR

A total of 1µg of RNA was reverse-transcribed for each condition. A transcript with accession number AY068123 was used as the internal control. FastSYBR Green Master Mix from Applied Biosystems was used in the StepOne Plus Real-Time StepOne Software. Triplicates were performed for each condition in the experiments and repeated two times independently of each other.

### RNAi experiments

The RNAi was facilitated via microinjections of dsRNA synthesized *in-vitro* as previously described^36^. Three pulses (containing 32 nL of dsRNA each) were given in the pre-pharyngeal area. Five injections were given in the span of twenty days. Control planarians were subjected to RNAi using a similar protocol but with a gene absent in the planarian genome. At twenty days post-first RNAi, animals were evaluated for macroscopic and microscopic manifestations.

### In-situ hybridization and Immunohistochemistry

Whole-mount *in-situ hybridization* was performed as previously described^37^. Immunohistochemistry was performed in Carnoy’s solution-fixed animals. Tissues in fixed samples were blocked for 4 hours and then incubated using primary antibodies overnight: αH3P 1:250 (Millipore Cat# 05-817R); 6G10 1:500 (DSHB Cat# 6G10-2C7); RAD51 1:500 (Abcam Cat# ab109107); and γH2AX antibody 1:1000 (ThermoFisher Cat# LF-PA0025). Samples were washed 6 x 10 mins in PBSTB and incubated with secondary antibody overnight: 1:500 HRP-conjugated Goat anti-rabbit antibody (Millipore Cat# 12-348) followed by washes 6 x 10 with PBSTB and mounted in VECTASHIELD mounting media. Samples were imaged with the Nikon AZ-100 microscope or the NIS Elements AR 3.2 software.

### Planarian tissue fixation

The samples for immunohistochemistry were euthanized with hydrochloric acid, and fixed with Carnoy’s solution, followed by hydrogen peroxide bleaching. Samples for whole-mount in situ hybridization (WISH) were euthanized with 5% N-acetyl cysteine, followed by a formaldehyde fixation, followed by a formamide/hydrogen peroxide bleaching.

### Growth Analysis

Animals were photographed every five days for a month using a Nikon AZ-100 multizoom microscope. The animal surface area was measured using ImageJ (https://imagej.net/ij/). At every time point, the fold change was compared to the control group.

### TUNEL

TUNEL assay was performed as described before using the ApopTag Red *in-situ* apoptosis detection kit^38^. Animals were fixed with NAC and formaldehyde. Samples were bleached overnight with PBSTx. The next day animals were rinsed with 1x PBS and incubated with TDT enzyme for 4 hours. After incubation, animals were rinsed with 50µl stop solution (provided in the kit), followed by a rinse in 1xPBS, and overnight incubation with DIG-POD diluted in blocking solution (provided in the kit) at 4°C. Following secondary incubation, animals were washed with PBSTB seven times every 15 minutes, followed by mounting and imaging. TUNEL^+^ cells were counted using Image J. Samples were imaged with a Zeiss LSM 880 confocal microscope.

### Comet

The assay was performed as previously described^39^. Briefly, cells were mechanically dissociated and suspended in calcium-magnesium-free (CMF) media. Cells strained and left at 37°C for 2 hours. Mixed with low melting point agarose and embedded on an agarose-covered glass slide. After the agarose solidifies, the coverslip is removed and placed in a Coplin jar with a lysing solution and left overnight. The next day, slides were placed in a neutralization buffer, followed by an electrophoresis buffer, and electrophoresis was done for 30 minutes at 12V, 300mA. The slides were removed from the electrophoresis buffer and washed with neutralization buffer. The neutralization buffer is then removed and placed in pre-chilled ethanol for 5 minutes at -20°C. The slides were then left to air dry for 1 hour and stained with Syber gold. Images were taken with the Nikon AZ-100 microscope and the NIS Elements AR 3.2 software. Comet tails were measured using Image-J as previously described^39^.

### Statistical Analysis

Data are expressed as the mean, standard error of the mean (SEM), or fold change SEM. Statistical analyses were performed in Prism, GraphPad software Inc.

## RESULTS

### *Dmap1* homolog is evolutionarily conserved in planarians

Previous research found that the *Schmidtea mediterranea* genome (Smed genome) lacks cytosine-dependent methylation and absence of DNMT1^39^. DMAP1 is a part of the DNMT1 multiprotein complex that facilitates DNA methylation, and it is also known to associate with other proteins implicated in acetylation and deacetylation^11–15,18,19,23–30,45–47^. We performed blast analyses using the publicly available database Planmine (https://planmine.mpinat.mpg.de/planmine/begin.do)^43^ and identified one *Dmap1* homolog (*Smed-Dmap1*) in the Smed genome. The human DMAP1 protein contains two domains, a SANT and a DMAP1 domain. We also found in the NCBI Conserved Domain Search (https://www.ncbi.nlm.nih.gov/Structure/cdd/wrpsb.cgi) that both the SANT and DMAP1 domains were present in *Smed-Dmap1* (Figure 1 A, Supplemental Figure S1). A protein alignment of DMAP1 between humans and *S. mediterranea* sequences revealed that DMAP1 proteins are 37% identical. The planarian SANT domain is 50% identical, and the DMAP1 domain is 32% identical to their human counterpart, respectively (Figure 1A). Overall, this suggests that the *Dmap1* sequence is evolutionarily conserved in *S. mediterranea*.

**Figure 1.**
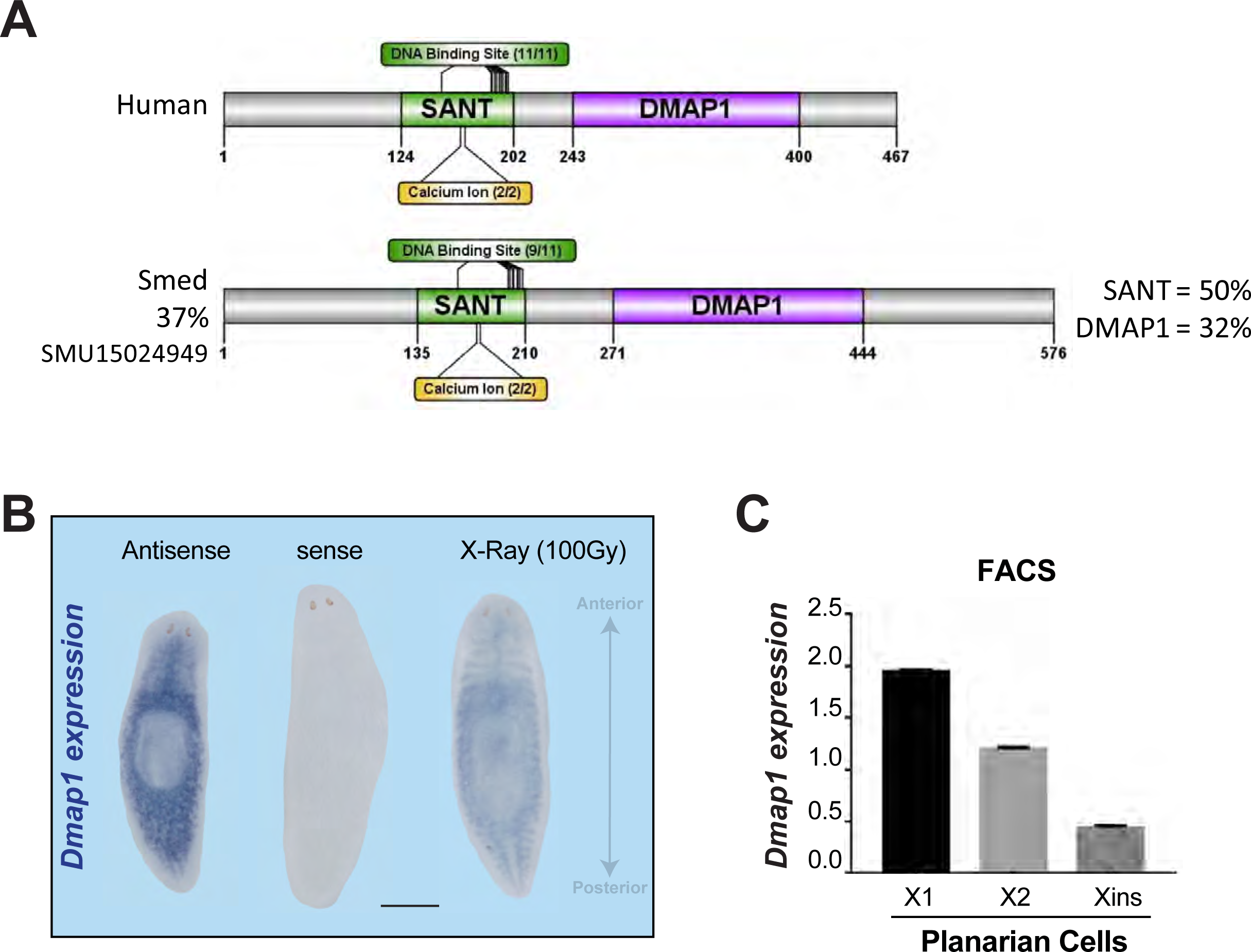
*Dmap1* is evolutionarily conserved and ubiquitously expressed in *S. mediterranea*. **A)** Protein conservation modeling of smed-DMAP1 relative to the human counterpart. Signature domains of DMAP1 include SANT and -DMAP1 with 50% and 32% homology. B) Whole mount in situ hybridization of Dmap1 with sense and antisense probes after a lethal dose of gamma radiation. C) *Dmap1* gene expression in planarians FACS-isolated cells. The planarian cells are irradiation-sensitive (X1 and X2) and insensitive (Xins) cells. Scale bar 400µm.

### *Dmap1* is ubiquitously expressed in planarians

To identify the spatial distribution and cell types expressing *Dmap1,* we performed whole-mount *in-situ* hybridization, flow cytometry, and *in-silico* analysis (Figures 1B, C, and Supplemental Figure S2). First, whole-mount *in-situ* hybridization revealed that *Dmap1* is ubiquitously expressed in the planarian body and appears enriched in the intestinal and parenchymal cells surrounding the pharynx (Figure 1B). We also confirmed the specificity of the signal by using a sense riboprobe that did not hybridize throughout the animal (Figure 1B). To identify whether the broad distribution of *Dmap1* expression involved stem cells and differentiated cells, we exposed planarians to lethal doses of gamma radiation (100 Gy) that are known to eliminate neoblasts (i.e., the planarian stem cells). The results show that *Dmap1* expression is reduced after the treatment with gamma radiation, suggesting its expression is present in both neoblasts and post-mitotic cells (Figure 1B). We also used flow cytometry to sort planarian cells and performed qPCR on three populations based on their sensitivity to ionizing radiation (Figure 1C). We found *Dmap1* expression throughout all three populations, including proliferating cells (X1) and non-proliferating cells (X2 and Xins). These results suggest that *Dmap1* is present in neoblasts and differentiated cells.

Further gene expression studies using single-cell databases were used to gain additional details of the cell types expressing *Dmap1* (Supplemental Figure S2). Planosphere (https://planosphere.stowers.org/), a public database, allowed us to visualize *Dmap1* expression in neoblast subsets and early lineage progenitors. These analyses revealed that most *Dmap1* expression was found in neoblast and early lineage progenitors (Supplemental Figure S2A-F). This finding is consistent with our results using gamma radiation that dramatically suppresses *Dmap1* expression (Figure 1B and Supplemental Figure S2A-C). We expanded the *in silico* analysis using another public database (Digiworm, https://digiworm.wi.mit.edu/) and found that trace amounts of *Dmap1* expression may be present in differentiated cells (Supplemental Figure 2G). Additionally, the predictive *Dmap1* expression pattern along the anteroposterior axis was inferred using Planmine (https://planmine.mpinat.mpg.de/planmine/begin.do). The results are consistent with the spatial distribution obtained in our *in-situ* hybridization pattern whereby *Dmap1* expression is more intense around the pre-pharyngeal and pharyngeal area, gradually decreasing towards the most posterior parts of the animal (Supplemental Figure 2H). Altogether, our findings reveal that *Dmap1* is broadly expressed throughout the planarian body with enrichment in neoblasts subpopulations.

### *Dmap1* is required for tissue maintenance in planarians

Next, we sought to functionally characterize *Dmap1* using RNA-interference (RNAi). We devised an RNAi strategy that consisted of five injections over 20 days (Figure 2A). We confirmed that this RNAi schedule effectively downregulates *Dmap1* expression by performing qPCR at the end of the injection time course (Figure 2B). Downregulating *Dmap1* resulted in macroscopic abnormalities after the fourth injection, and over 90% of the animals presented tissue alterations by 20 days post-RNAi (Figure 2C). The tissue abnormalities included tissue degeneration involving head regression and tissue rippling (49%) and lesions in the posterior parts of the animals, including tail loss (45%) that, in all cases, led to lethality. The tissue degeneration was related to missing areas in the anterior, also known as head regression, while the lesions in the posterior tissues surrounded the pharynx area (Figure 2C). We observed that over the course of the *Dmap1* phenotype, the experimental group showed a continuous decrease in size, reaching about 50% reduction in length by 15 days after the first dsRNA injection. The group continued to decrease to a quarter of the original size after 25 days (Figure 2D). Significantly, about 50% of the *Dmap1(RNAi)* animals died within the first 10 days post-RNAi; by day 30, all animals were dead (Figure 2E).

**Figure 2.**
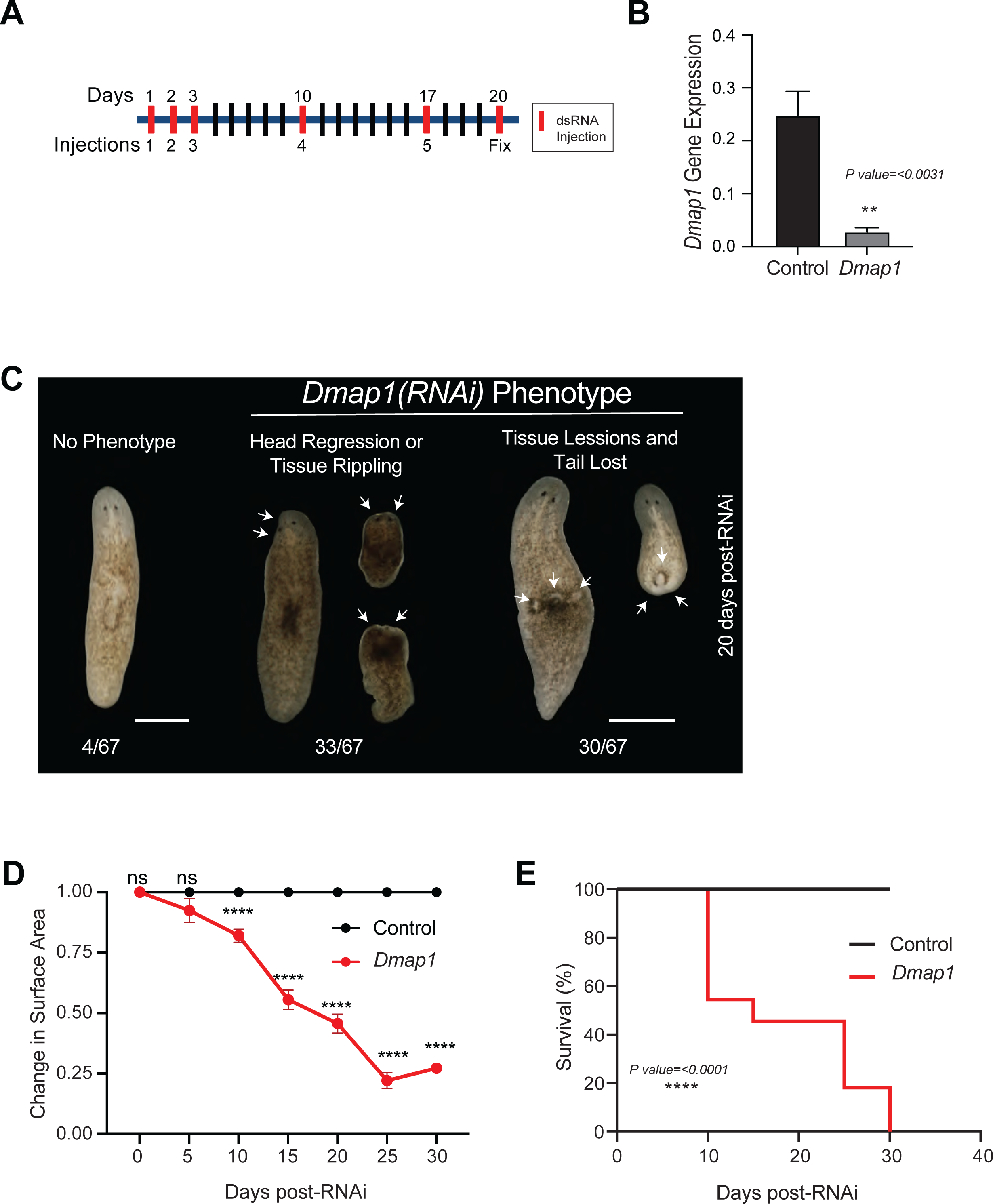
*Dmap1* is essential for tissue maintenance and survival. **A)** RNAi schedule facilitated by *Dmap1* dsRNA microinjections. Five injections (red bars) were administered in 20 days before fixation. **B)** *Dmap1* expression levels were measured with qPCR using whole animals at 20 days post-RNAi. Gene expression experiments were performed in triplicates with ten samples each; **p<0.0031; Unpaired t-test. **C)** Representative live images of *Dmap1* RNAi animals 20 days post-RNAi. Arrows indicate different abnormalities, including head regression or tissue rippling, tissue lesions, and tail loss in *Dmap1(RNAi)* animals. The numbers at the bottom represent the animals in each category. **D, E)** Change in surface and Kaplan-Meier probability of survival over 30 days; ****<0.0001; two-way ANOVA. The experiments in **C-E** consisted of seven biological replicates with about ten animals each. Scale bar 200µm.

### *Dmap1* regulates neoblast division and regeneration

Neoblasts are the only cells capable of dividing in planarians, and they constantly proliferate to support tissue turnover and regeneration of damaged parts. We hypothesized that macroscopic epithelial tissue lesions observed in *Dmap1(RNAi)* may result from neoblast proliferation and differentiation defects. First, we analyzed the neoblast division with immunohistochemistry using an anti-phosphorylated histone-3 (anti-H3P) antibody that labels mitotic cells (Figures 3A, B). We analyzed four time points every five days to learn about potential changes in neoblast proliferation (5, 10, 15, 20). These experiments revealed that after five days of RNAi, there was a 20% increase in neoblast division in the experimental group. However, over the remaining 15 days, levels of cell division were similar between the control and the experimental group (Figures 3A, B).

**Figure 3.**
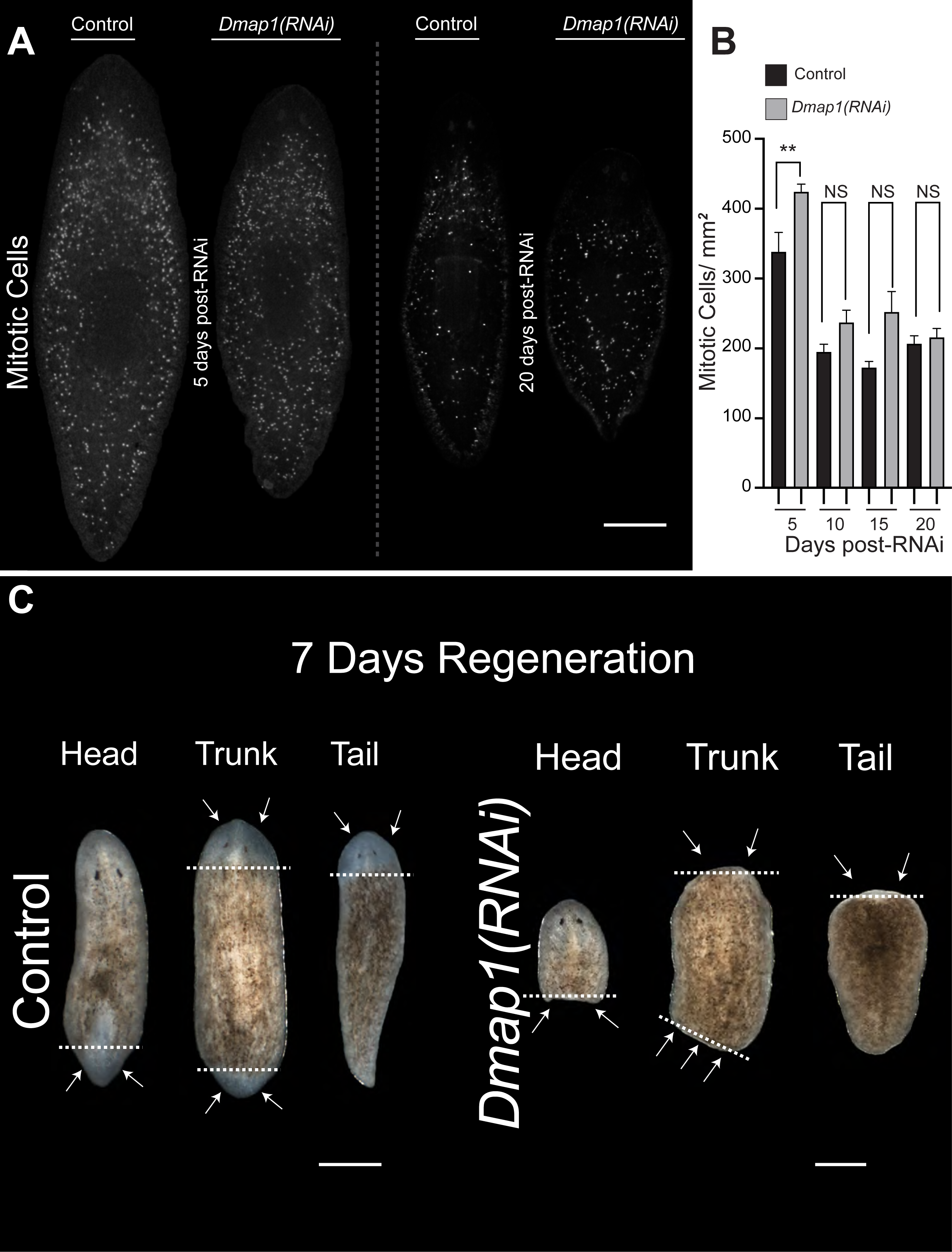
*Dmap1* regulates cell division and tissue regeneration. **A)** Spatial distribution of mitotic activity in whole mount immunostaining against phospho-histone H3 (Ser10) (H3P) at 5- and 20-days post-RNAi. **B)** Levels of mitoses in a 20-day time course; **p<0.0046; one-way ANOVA. **C)** Representative live images of regenerating fragments (head, trunk, and tail) of control and *Dmap1(RNAi)* animals post-amputation. White dashes indicate the area of amputation. White arrows show the location of newly regenerated tissues (control) or the failure to regenerate in all animals of the experimental group. A-C consisted of five biological replicates involving 8-52 samples per time point. Scale bar 200µm.

To further investigate the effects of *Dmap1(RNAi)* on neoblast function, the regeneration response was evaluated. Neoblasts are necessary to repair missing parts upon amputation. Therefore, we amputated planarians after the fourth dsRNA injection (i.e., day 11) and followed for a week the regeneration of heads, trunks, and tail fragments (Figure 3C). The results demonstrated that while control fragments regenerated missing parts, the totality of *Dmap1(RNAi)* animals could close wounds but failed to form blastemas and repair missing parts (Figure 3C). The lack of regeneration in the presence of mitotic neoblasts in the experimental group suggests that *Dmap1* is required for the proper neoblast response to injury.

### *Dmap1* is required for neoblast differentiation

Since epithelial lesions characterize the macroscopic defects in the *Dmap1(RNAi)* phenotype, we decided to investigate whether the transition from neoblasts to epithelial lineage was altered. Experiments using fluorescent whole-mount *in-situ* hybridization (FISH) with markers of neoblasts *(piwi)*, early (*prog2)*, and late stages of epithelial differentiation (*agat, ifb,* and *vim1*) revealed a marked reduction in expression in the experimental group at 20 days post-RNAi (Figure 4A). Between 80-100% of the experimental animals displayed reduced expression of the neoblast and the epithelial markers, suggesting *Dmap1* is a critical regulator of neoblast function and differentiation toward the epithelial fate. The reduction of expression in the experimental group was more notorious in the anterior part of the animal (Figure 4B). These findings are consistent with other phenotypes involving dysfunctional neoblasts^40^. In some cases, signal discontinuity of differentiated epithelial cells (i.e., *ifb*) suggested a lack of renewal or unrepaired epithelial damage.

**Figure 4.**
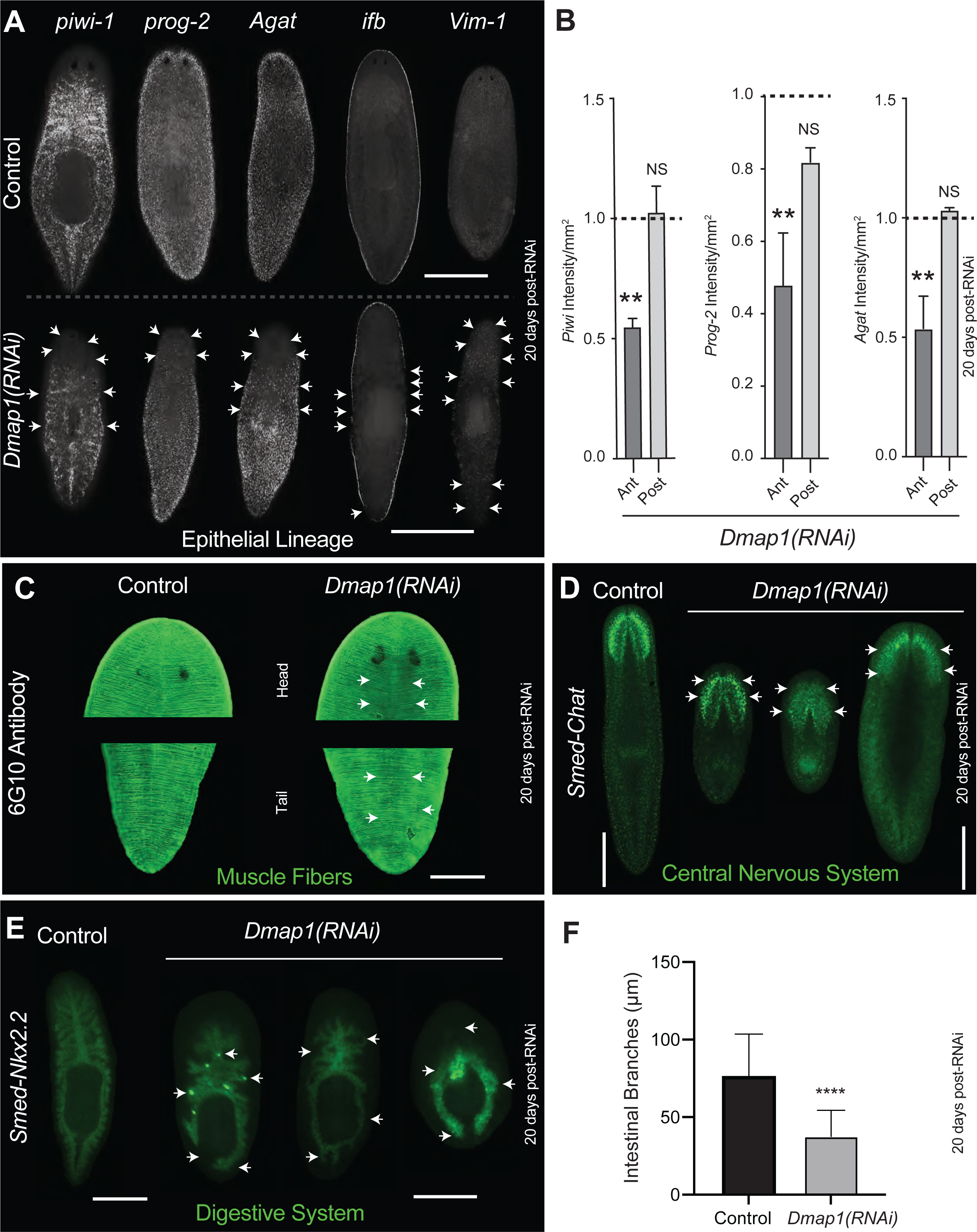
*Dmap1* is required for neoblast differentiation. **A)** Fluorescent *in situ* hybridization of markers associated with the epithelial lineage in planarians. These include the pan-neoblast marker (*piwi-1*), early epidermal progenitor marker (*prog-2*), late epidermal progenitor marker (*agat*), and differentiated epidermal markers (*ifb* and *vim-1*) in control and *Dmap* RNAi animals 20 days post RNAi. **B)** Changes in intensity expression per area and the anterior and posterior region of *Dmap1(RNAi)* animals (control values are represented with a dashed line). Reliable changes were obtained for early differentiation markers at 20 days; **p<0.0039; one-way ANOVA. **C)** Whole mount immunostaining using the muscle marker 6G10 antibody. **D-E)** Fluorescent *in situ* hybridization using a neural marker (*Smed-Chat*) and intestinal marker (*Smed-Nkx2.2*), respectively. Arrows depicting *Dmap-1(RNAi)*-induced abnormalities in the expression of muscle fibers and nervous and intestinal tissues. **F)** Secondary intestinal branch length measurements of control and *Dmap1(RNAi)* animals. The measurements were restricted to the pre-pharyngeal area due to a lack of resolution in the areas below the pharynx. Two biological replicates with four animals each, ****p<0.0001 Unpaired t-test with Welch’s correction. **A, D, E** Two-three biological replicates with about five to ten animals each. Scale bar 200µm.

We also explored whether *Dmap1(RNAi)* affects the neoblast potential to differentiate into other lineages (Figures 4C-F). First, qualitative observation of muscle cells with the antibody 6G10 (Ross et al., 2015), evidenced areas with low muscle presence and disorganization of the muscle fibers (Figure 4C). We also found that experimental animals displayed a reduced expression throughout the body for the neural marker *Smed-Chat* (Figure 4D). The *in situ* hybridization with the intestinal marker *Nkx2.2* showed most animals with disorganized intestinal branches and variations in the marker expression levels (Figure 4E). Quantification of the intestinal branches in the anterior part of the animal (i.e., head and pre-pharyngeal regions) evidenced a 43% reduction in the number of branches (14 vs 6 branches in control and experimental animals, respectively). This was also accompanied by a 39% reduction in the length of the intestinal secondary branches (Figure 4F). Altogether, the results suggest that *Dmap1* is essential for neoblast differentiation across different lineages.

### *Dmap1* regulates cell death fate in the presence of DNA damage

Although the late stages of the *Dmap1* phenotype showed increased epithelial lesions, it is accompanied by similar levels of neoblast division between the control and experimental groups. Next, we performed analyses of cell death with whole-mount immunostaining using TUNEL (Figure 5A). Cell death is always present, and it is necessary to remove old or damaged cells during the process of cellular turnover. Thus, animals in the control group display sparse levels of cell death throughout their body. However, we observed that at 20 days after *Dmap1(RNAi)*, there was a seeming increase in the density of the TUNEL signal around the epithelial lesions (arrows in Figure 5A). The increase in TUNEL signal was commonly seen as clusters of cells clumped around the damaged area. The confocal analyses demonstrated that clustered cell death was distributed across different layers along the dorsoventral axis. These results suggested that *Dmap1(RNAi)-*induced cell death is widely distributed across different cellular layers and likely involving different cell types.

**Figure 5.**
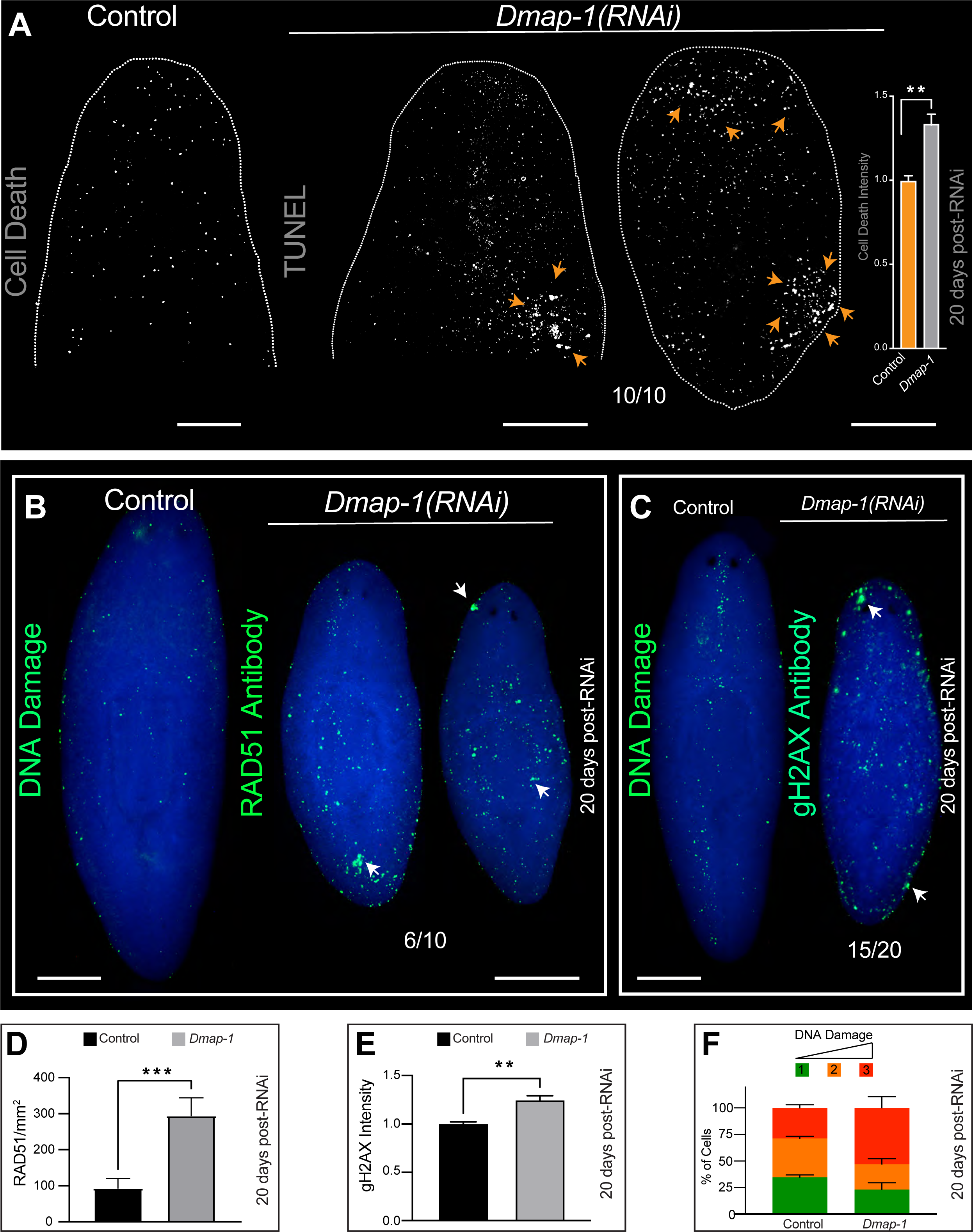
*Dmap1* regulates cell death and DNA integrity. **A)** Spatial distribution of cell death using TUNEL staining (white dots). Arrows show examples where TUNEL^+^ cells clustered. Samples on the left show higher magnification of the anterior planarian region, and the image on the right displays a whole animal subjected to *Dmap1(RNAi)*. The graph on the right displays the levels of intensity of Tunel positive cells in different segments of the samples. **B, C)** Whole mount immunostaining against RAD51, and gH2AX antibodies (green signal). Arrows indicate places with clustered signal likely associated with DNA damage. **D, E)** Levels of RAD51 and gH2AX^+^ foci 20 days post-RNAi. **F)** DNA integrity measured with the COMET assay. Color-coded categorization of DNA damage showing (1) green, no DNA damage, (2, 3) mild and severe DNA damage (orange and red respectively). Three biological replicates with about ten animals each. **B-E)** Two biological replicates with 5-10 animals each; **p<0.065; ***p<0.001, unpaired t-test with Welch’s correction. Scale bar 200µm.

The *Dmap1* function in other organisms has been associated with the maintenance of DNA stability and repair^3,19–22^. Therefore, we analyzed the DNA damage response and DNA repair with immunostaining using the RAD51 and gamma-H2AX (gH2AX) antibodies and the Comet assay. The phosphorylation of the histone H2aX protein is one of the earliest manifestations of DNA damage, and the RAD51 protein is essential for the repair of DNA double-strand breaks; in planarians, its expression is enriched in the neoblasts^41–44^. Our analyses identified a dramatic increase in both gH2AX and RAD51 signals across the planarian’s body of experimental animals (Figure 5B-E). We also noticed cell clusters associated with tissue lesions. Furthermore, we also detected that not only did the activation of the DNA damage response occur in the experimental animals but also a ∼25% increase in DNA double-strand breaks, as evidenced using the Comet assay, occurred at 20 days post-*Dmap1(RNAi)* (Figure 5F). These experiments demonstrated that *Dmap1* is essential for DNA damage repair in planarians.

### *Dmap1* regulates the expression of polarity markers in planarians

DMAP1 is known to regulate the activation and repression of transcription. To test whether DMAP1 function is evolutionarily conserved in planarians, we analyzed the expression pattern of genes associated with axial polarity determination, whose transcription is spatially regulated by unknown factors. Inhibitors and activators of the Wnt signaling mediate the anteroposterior axial polarity in planarians. *Notum* and *wnt11-2* are Wnt mediators specifically expressed by a few muscle cells at either the anterior or posterior pole in planarians. Our whole-mount fluorescent *in situ* hybridization (WISH) experiments confirmed the expression of *notum* and *wnt11-2* at the anterior and posterior poles of the control animals (Figure 6A, C). Strikingly, the expression pattern of AP polarity markers in *Dmap1(RNAi)* animals was ectopically found across the body. To illustrate these findings, we divided into four sections along the AP axis and recorded the location of the expression. Notum in the control is expressed only in section one (i.e., at the anterior tip, in front of the eyes). Nonetheless, we found that 100% of *Dmap1(RNAi)* animals at 20 days showed notum expression in segments 1 and 2 and about 75% in segments 3 and 4 (Figure 6B). The posterior marker *wnt11-2* was also found ectopically expressed in segments 1-3 and in clusters, suggesting not only alterations in spatial distribution and number of cells and intensity (Figure 6C, D).

**Figure 6.**
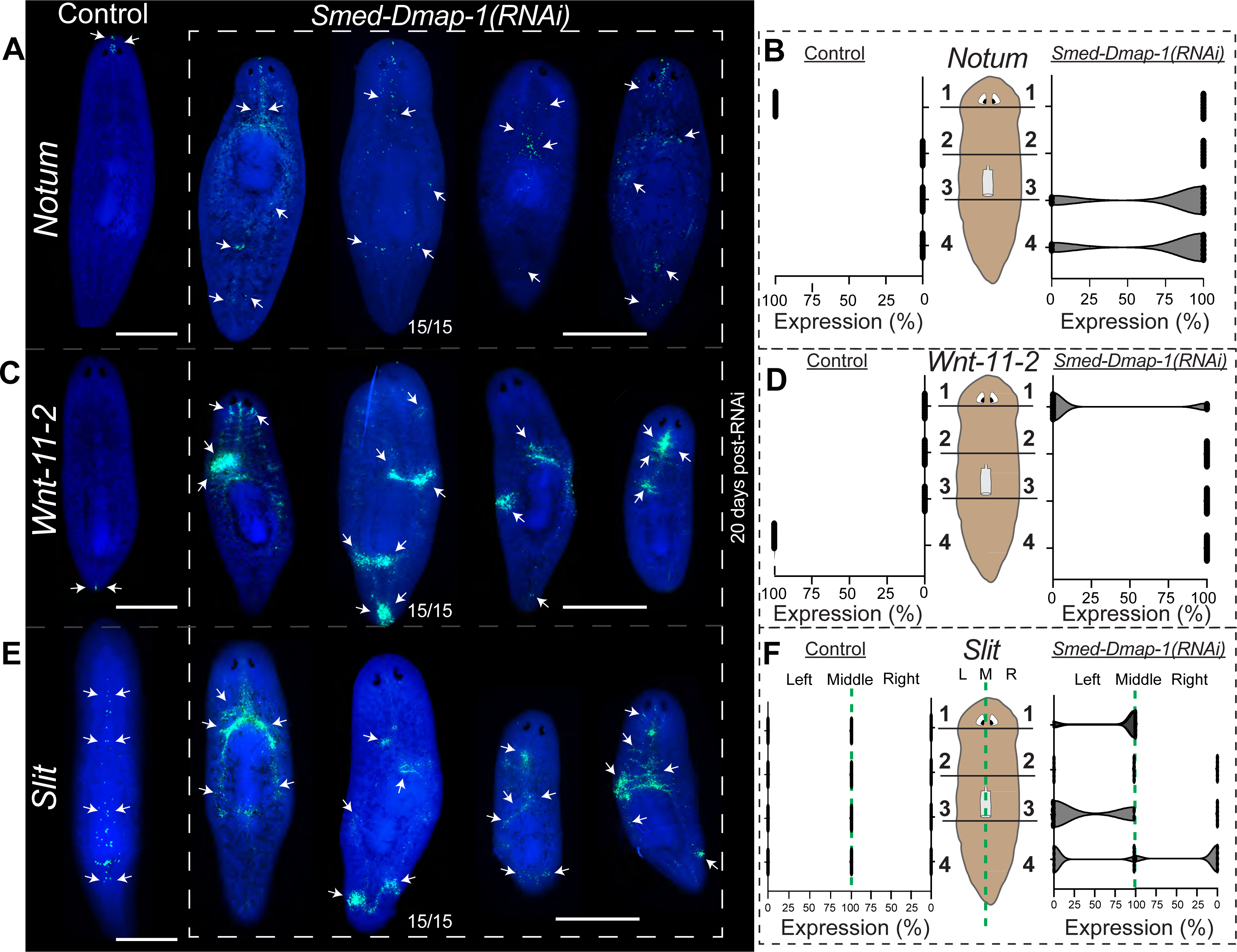
*Dmap1* restricts the expression of axial polarity markers. **A)** Fluorescent *in situ* hybridization (FISH) using a probe against anterior marker (*Smed-notum*), **C)** Posterior marker (*Smed-Wnt-11-2*), and **E)** midline marker (*Smed-Slit*). Arrows depict normal or ectopic gene expression in control and experimental animals (15/15), respectively. **B, D, F)** Violin plots representing the percentage of gene expression along the anteroposterior and mediolateral axis. Animals were divided into four regions, starting with the head and ending with the tail region (1-4). In addition, the *Smed-Slit* signal was evaluated on the left and right sides of the animals (the green dotted line illustrates the middle axis). Three biological replicates with 5 animals each. Scale bar 200µm.

Furthermore, we also analyzed the mediolateral axis marker *slit*, which is expressed by a restricted group of cells in the midline from anterior to posterior. The *slit* expression in *Dmap1(RNAi)* was observed outside of the midline and with inconsistent patterns and expression intensities across all segments (Figure 6E, F). Together, the results show that *Dmap1* is essential to maintain restricted expression patterns along the AP and mediolateral axis.

## Discussion

Our work examined the role of the planarian homolog *Dmap1* during systemic cellular turnover and regeneration. *Dmap1* is essential for planarian tissue maintenance, regeneration, and survival. The data show that *Dmap1* is critical for the differentiation of neoblasts into different lineages, maintenance of DNA integrity, and necessary for the spatial expression of axial polarity markers.

Previous studies suggested that DMAP1 functions independently and as a subunit of distinct complexes involved in the repression or activation of transcription^8,15,16,23,45^. DMAP1 consists of two protein domains: one SANT domain, which mediates histone tail binding, and a DMAP1 domain, a methylase that represses transcription^8,45^. The human genome contains one DMAP1. The two domains present in DMAP1 are highly conserved throughout the vertebrates. In *S*. *mediterranea,* we found one DMAP1 orthologous to vertebrates containing both the SANT and DMAP1 domains with a 37.4% identity to DMAP1 in humans. Like its human counterpart, the planarian DMAP1 is ubiquitously expressed^46^. Our functional analyses suggest that *Dmap1* may act as a transcriptional repressor. We observed that when the *Dmap1* function is abrogated, some cells gain the capacity to ectopically express genes with different intensities. These results suggest that *Dmap1* may be crucial in regulating the location and levels of transcription in adult tissues (see below).

Functional disruption of DMAP1 is lethal in insects and mammals^23,47,48^. Our results demonstrate that *Dmap1* is necessary for planarian survival, arguing for evolutionary conservation at the organismal level in adult stages. Recent studies showed that DMAP1 facilitates tissue maintenance and commitment to differentiation in embryonic stem cells^15,20,22,23^. We also found that *Dmap1(RNAi)* leads to a reduction in expression of the neoblast marker *piwi-1,* and early and late epithelial progenitors suggest that *Dmap1* contributes to epithelial lineage differentiation, affecting epithelial cell turnover and overall maintenance. This function was also extended to the nervous and intestinal systems and muscle cells. Although our findings suggest a generalized lack of tissue maintenance, the defects were not distributed at the same levels across tissues, which may argue for varying degrees of *Dmap1* regulation in neoblast differentiation. Intriguingly, despite high levels of DNA damage, the neoblast division is not reduced, but their function to attend to and repair tissues is compromised. These findings strongly suggest that *Dmap1* is an essential component of the neoblast function to repair tissues. Together, our results show the functional conservation of *Dmap1* in regulating stem cell biology.

The functional disruption of *Dmap1* in planarians led to increased gH2AX and RAD51 signals and extensive DNA damage. These findings are consistent with an evolutionary role for DMAP in sensing DNA damage and repair, which has been demonstrated in hematopoietic stem cells and other mammalian cells^19–21,45^. The mechanisms of how downregulation of *Dmap1* leads to DNA damage in planarians need to be further investigated. Nonetheless, since the DNA repair machinery is highly conserved in *S. mediterranea*, it is possible that disruption of ATM signaling and/or its association with DNMT1 may contribute, as previously demonstrated in mammals^19,21^. Additionally, it is possible that the planarian *Dmap1* associates with other complexes, such as NuA4 histone acetyltransferase complex that acetylates histone-4 and histone 2A, which is essential for transcriptional regulation, chromosomal stability, and homologous recombination repair^13^. High levels of DNA damage in the *Dmap1* phenotype may result in increased levels of cell death throughout the planarian body. Although, this was the case, we often observed cell death clustered in areas where tissue damage occurred. We are prompted to speculate that *Dmap1* contributes to cell death fate decisions in the presence of DNA damage in areas of tissue injury that are known to require stereotypic patterns of apoptosis^49^. Additional experiments are needed to resolve whether neoblasts divide with DNA damage in the *Dmap1* phenotype.

An advancement in the tools available in the planarian field has made it possible to interrogate the roles that epigenetics has in regulating planarian neoblast^29,33^. Previous studies have shown that loss of function of histone methyltransferases and loss of function of the methyl-CpG Binding Domain 2/3 (*mbd2/3*) lead to an impairment of regeneration, stem cell depletion, and differentiation^29,34,50^. Implementing the ChIP-seq protocol in planaria provided insight into the evolution of bivalent promoters that function as activators and repressors^29,32^. Planarians are among the earliest organisms to show epigenetic regulation for adult stem cell-mediated cell turnover and regeneration^32,51–53^. Additional studies in planarians have shown that MLL3/4, another histone methyltransferase, regulates oncogenes and tumor suppressors^25,29^. *Dmap1* appears to complement some of these functions as it is necessary for proper neoblast differentiation and tissue repair. We found it intriguing that not all tissues have similar levels of defects after *Dmap1(RNAi)*. For example, muscle and the nervous system were not as impacted as the epithelial cells and the intestinal compartment. This may imply that *Dmap1* function differs across tissues, but additional experiments are needed to resolve the relevance of *Dmap1* in individual cell lineages.

Uniquely, our findings implicate *Dmap1* in regulating gene expression of axial polarity markers. On one side, the planarian *Dmap1* shows functional conservation as a transcription repressor. Once the *Dmap1* function is silenced, some cells acquire the capacity to ectopically express positional control genes (PCGs) normally transcribed by muscle cells. It remains unclear whether the ectopic expression of PCGs may induce new axial polarity in the animal. We did not observe ectopic organ or behavioral changes before animals died. Likewise, the number of cells and the intensity in which these polarity markers are expressed was abnormal, suggesting *Dmap1* also regulates spatial expression, the level and number of cells associated with AP, and the mediolateral axis. The mechanisms linking *Dmap1* with Wnt signaling are uncertain, but there is evidence that DNMT1 interacts with β-catenin to regulate Wnt signaling^10^. However, a previous work in *S. mediterranea* did not find any DNMTs^29^. It is possible that DMAP1 may interact independently with β-catenin to regulate Wnt signaling, but further studies need to be done in order to verify this interaction. Overall, this finding presents *Dmap1* as a novel regulator that restricts positional control genes to specific regions in the planarian body, and loss of function of *Dmap1* leads to ectopic expression of positional control genes.

In conclusion, DNA methyltransferase Associated Protein has multiple roles in planaria, some conserved and some novel. Our results present that *Dmap1* engages in maintenance, survival, neoblast proliferation, neoblast differentiation, homologous recombination, and maintenance of polarity markers. Further investigations are necessary to delineate molecular mechanisms coordinating DMAP1 function in adult tissue turnover and regeneration.

## Acknowledgments

We thank members of the Oviedo lab for insightful discussions and comments on the manuscript. The 6G10 antibody was obtained from the Developmental Studies Hybridoma Bank, created by the NICHD of the NIH and maintained at the University of Iowa, Department of Biology.

## Author contributions

S.R., P.G.B., P.K., and N.J.O. conceived, designed, and executed the experiments, and S.R. and N.J.O. wrote the manuscript. All authors read the manuscript, provided comments, and approved the final version.

## Figure Legends

**Supplemental Figure 1.**
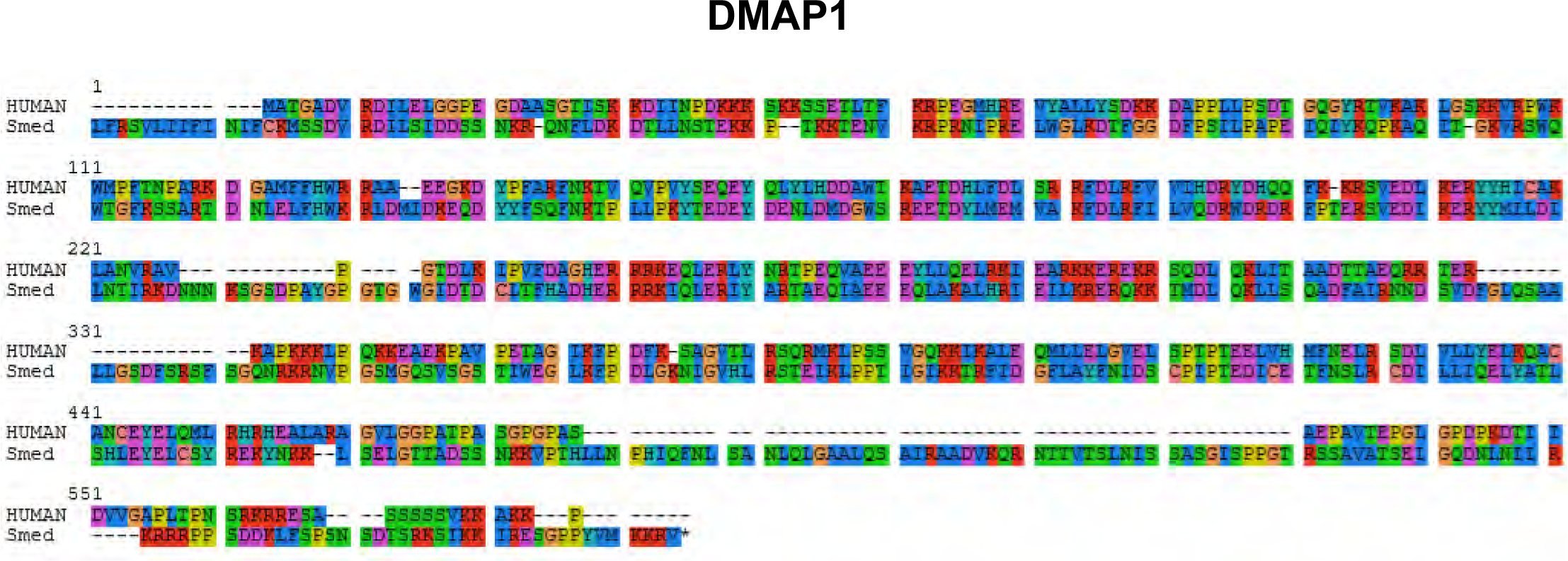
DMAP1 protein sequence conservation between humans and *S. mediterranea*. Protein alignment sequence generated by CLUSTALW between humans and *S. mediterranea* DMAP1 predicts 32% conservation with its human counterpart. The color-coded amino acids display hydrophobic (blue), positive charge (red), negative charge (magenta), polar (green), cysteines (pink), glycines (orange), prolines (yellow), aromatic (cyan), and unconserved (white).

**Supplemental Figure 2.**
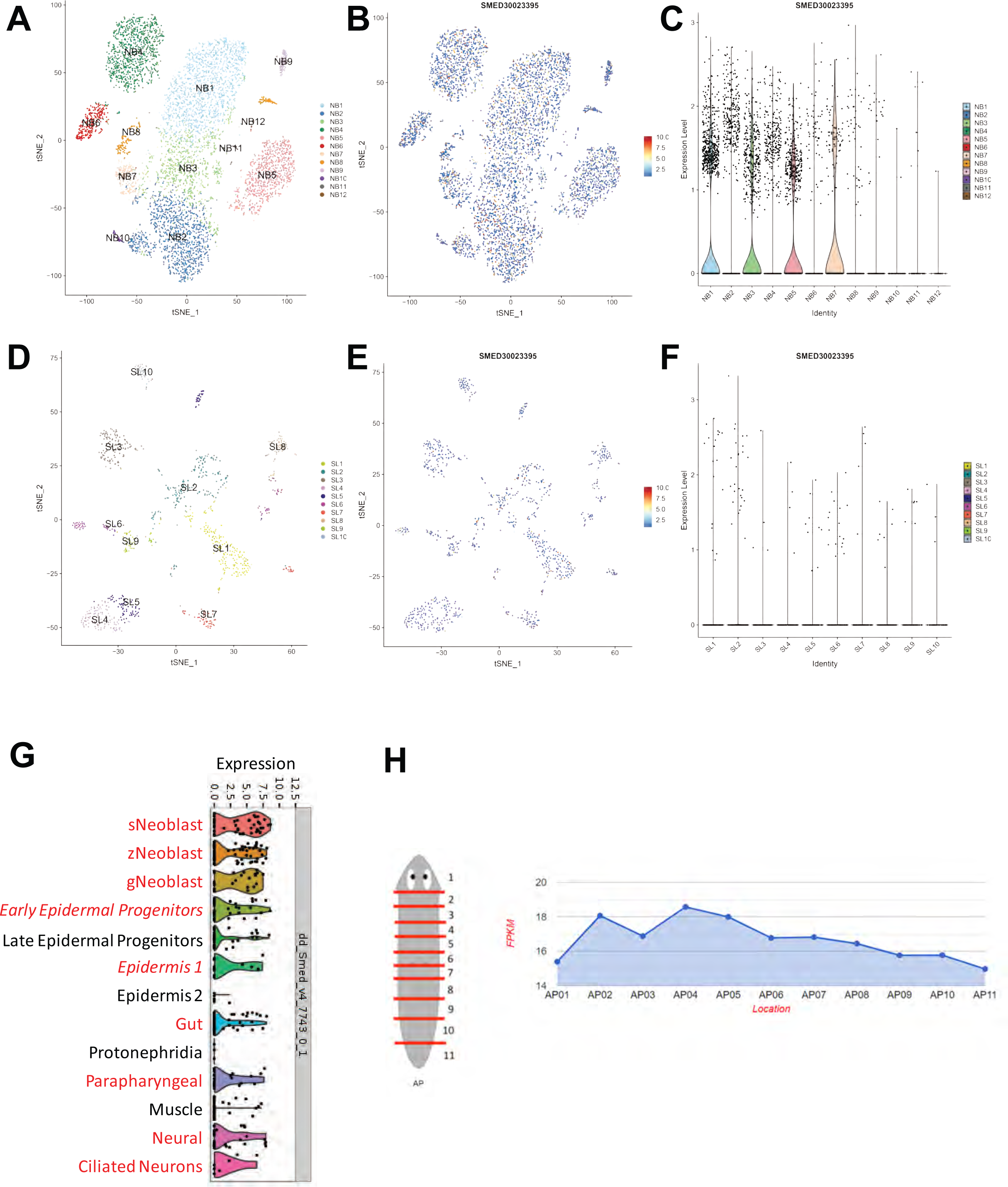
D*m*ap1 is ubiquitously expressed in planarians. **A-H)** *in-silico* analysis of Dmap1 expression in neoblasts, differentiated cells, and along the anteroposterior (AP) axis. **A-F)** shows *Dmap1* predicted expression across twelve neoblast subpopulations (colored dots and clusters). Most *Dmap1* expression is predicted to be in neoblast subtypes NB1, NB3, NB5, and NB7. G) the predicted Dmap1 expression in three neoblast subpopulations (sigma, zeta, and gamma) and neoblast progenitors and differentiated cells. Red highlighted cell types express higher levels of *Dmap1*. H) *Dmap1* expression levels along the AP axis divided in eleven segments from the anterior to the posterior. The *in-silico* analyses were generated using public databases including Planosphere, Digiworm, and Planmine.

## References

1. Moore, L.D., Le, T., and Fan, G. (2013). DNA methylation and its basic function. Neuropsychopharmacology 38, 23–38. 10.1038/npp.2012.112.

2. He, S., and Feng, X. (2022). DNA methylation dynamics during germline development. J Integr Plant Biol 64, 2240–2251. 10.1111/jipb.13422.

3. Mattei, A.L., Bailly, N., and Meissner, A. (2022). DNA methylation: a historical perspective. Trends Genet 38, 676–707. 10.1016/j.tig.2022.03.010.

4. Borowczyk, E., Mohan, K. N., D’Aiuto, L., Chaillet, J. R. (2009). Identifiation of a region of the DNMT1 methyltransferase that regulates the maintenance of genomic imprints. PNAS 106.

5. Rountree, M.R., Bachman, K.E., and Baylin, S.B. (2000). DNMT1 binds HDAC2 and a new co-repressor, DMAP1, to form a complex at replication foci. Nat Genet 25, 269–277. 10.1038/77023.

6. Liu, Z., and Fisher, R.A. (2004). RGS6 interacts with DMAP1 and DNMT1 and inhibits DMAP1 transcriptional repressor activity. J Biol Chem 279, 14120–14128. 10.1074/jbc.M309547200.

7. Muromoto, R., Sugiyama, K., Takachi, A., Imoto, S., Sato, N., Yamamoto, T., Oritani, K., Shimoda, K., and Matsuda, T. (2004). Physical and functional interactions between Daxx and DNA methyltransferase 1-associated protein, DMAP1. J Immunol 172, 2985–2993. 10.4049/jimmunol.172.5.2985.

8. Kang, B.G., Shin, J.H., Yi, J.K., Kang, H.C., Lee, J.J., Heo, H.S., Chae, J.H., Shin, I., and Kim, C.G. (2007). Corepressor MMTR/DMAP1 is involved in both histone deacetylase 1- and TFIIH-mediated transcriptional repression. Mol Cell Biol 27, 3578–3588. 10.1128/MCB.01808-06.

9. Xiang, J., Luo, F., Chen, Y., Zhu, F., and Wang, J. (2014). si-DNMT1 restore tumor suppressor genes expression through the reversal of DNA hypermethylation in cholangiocarcinoma. Clin Res Hepatol Gastroenterol 38, 181–189. 10.1016/j.clinre.2013.11.004.

10. Song, J., Du, Z., Ravasz, M., Dong, B., Wang, Z., and Ewing, R.M. (2015). A Protein Interaction between beta-Catenin and Dnmt1 Regulates Wnt Signaling and DNA Methylation in Colorectal Cancer Cells. Mol Cancer Res 13, 969–981. 10.1158/1541-7786.MCR-13-0644.

11. Martisova, A., Holcakova, J., Izadi, N., Sebuyoya, R., Hrstka, R., and Bartosik, M. (2021). DNA Methylation in Solid Tumors: Functions and Methods of Detection. Int J Mol Sci 22. 10.3390/ijms22084247.

12. Doyon, Y., and Côté, J. (2004). The highly conserved and multifunctional NuA4 HAT complex. Curr Opin Genet Dev 14, 147–154. 10.1016/j.gde.2004.02.009.

13. Doyon, Y., Selleck, W., Lane, W.S., Tan, S., and Côté, J. (2004). Structural and functional conservation of the NuA4 histone acetyltransferase complex from yeast to humans. Mol Cell Biol 24, 1884–1896. 10.1128/mcb.24.5.1884-1896.2004.

14. Cai, Y., Jin, J., Florens, L., Swanson, S.K., Kusch, T., Li, B., Workman, J.L., Washburn, M.P., Conaway, R.C., and Conaway, J.W. (2005). The mammalian YL1 protein is a shared subunit of the TRRAP/TIP60 histone acetyltransferase and SRCAP complexes. J Biol Chem 280, 13665–13670. 10.1074/jbc.M500001200.

15. Gorrini, C., Squatrito, M., Luise, C., Syed, N., Perna, D., Wark, L., Martinato, F., Sardella, D., Verrecchia, A., Bennett, S., et al. (2007). Tip60 is a haplo-insufficient tumour suppressor required for an oncogene-induced DNA damage response. Nature 448, 1063–1067. 10.1038/nature06055.

16. Fazzio, T.G., Huff, J.T., and Panning, B. (2008). An RNAi screen of chromatin proteins identifies Tip60-p400 as a regulator of embryonic stem cell identity. Cell 134, 162–174. 10.1016/j.cell.2008.05.031.

17. Jacquet, K., Fradet-Turcotte, A., Avvakumov, N., Lambert, J.P., Roques, C., Pandita, R.K., Paquet, E., Herst, P., Gingras, A.C., Pandita, T.K., et al. (2016). The TIP60 Complex Regulates Bivalent Chromatin Recognition by 53BP1 through Direct H4K20me Binding and H2AK15 Acetylation. Mol Cell 62, 409–421. 10.1016/j.molcel.2016.03.031.

18. Devoucoux, M., Fort, V., Khelifi, G., Xu, J., Alerasool, N., Galloy, M., Wong, N., Bourriquen, G., Fradet-Turcotte, A., Taipale, M., et al. (2022). Oncogenic ZMYND11-MBTD1 fusion protein anchors the NuA4/TIP60 histone acetyltransferase complex to the coding region of active genes. Cell Rep 39, 110947. 10.1016/j.celrep.2022.110947.

19. Negishi, M., Chiba, T., Saraya, A., Miyagi, S., and Iwama, A. (2009). Dmap1 plays an essential role in the maintenance of genome integrity through the DNA repair process. Genes Cells 14, 1347–1357. 10.1111/j.1365-2443.2009.01352.x.

20. Koizumi, T., Negishi, M., Nakamura, S., Oguro, H., Satoh, K., Ichinose, M., and Iwama, A. (2010). Depletion of Dnmt1-associated protein 1 triggers DNA damage and compromises the proliferative capacity of hematopoietic stem cells. Int J Hematol 91, 611–619. 10.1007/s12185-010-0563-3.

21. Penicud, K., and Behrens, A. (2014). DMAP1 is an essential regulator of ATM activity and function. Oncogene 33, 525–531. 10.1038/onc.2012.597.

22. Lee, G.E., Kim, J.H., Taylor, M., and Muller, M.T. (2010). DNA methyltransferase 1-associated protein (DMAP1) is a co-repressor that stimulates DNA methylation globally and locally at sites of double strand break repair. J Biol Chem 285, 37630–37640. 10.1074/jbc.M110.148536.

23. Mohan, K.N., Ding, F., and Chaillet, J.R. (2011). Distinct roles of DMAP1 in mouse development. Mol Cell Biol 31, 1861–1869. 10.1128/MCB.01390-10.

24. Wang, K., Liang, Q., Li, X., Tsoi, H., Zhang, J., Wang, H., Go, M.Y., Chiu, P.W., Ng, E.K., Sung, J.J., and Yu, J. (2016). MDGA2 is a novel tumour suppressor cooperating with DMAP1 in gastric cancer and is associated with disease outcome. Gut 65, 1619–1631. 10.1136/gutjnl-2015-309276.

25. Mihaylova, Y., Abnave, P., Kao, D., Hughes, S., Lai, A., Jaber-Hijazi, F., Kosaka, N., and Aboobaker, A.A. (2018). Conservation of epigenetic regulation by the MLL3/4 tumour suppressor in planarian pluripotent stem cells. Nat Commun 9, 3633. 10.1038/s41467-018-06092-6.

26. Newmark, P.A., and Sanchez Alvarado, A. (2002). Not your father’s planarian: a classic model enters the era of functional genomics. Nat Rev Genet 3, 210–219. 10.1038/nrg759.

27. Reddien, P.W. (2018). The Cellular and Molecular Basis for Planarian Regeneration. Cell 175, 327–345. 10.1016/j.cell.2018.09.021.

28. Tweedie, S., Charlton, J., Clark, V., and Bird, A. (1997). Methylation of genomes and genes at the invertebrate-vertebrate boundary. Mol Cell Biol 17, 1469–1475. 10.1128/MCB.17.3.1469.

29. Dattani, A., Sridhar, D., Aboobaker, A. A. (2019). Planarian flatworms as a new model system for understanding epigenetic regulation of stem cell pluripotency and differentiation. Seminars in Cell & Developmental Biology 87.

30. Pascual-Carreras, E., Marin-Barba, M., Castillo-Lara, S., Coronel-Cordoba, P., Magri, M.S., Wheeler, G.N., Gomez-Skarmeta, J.L., Abril, J.F., Salo, E., and Adell, T. (2023). Wnt/beta-catenin signalling is required for pole-specific chromatin remodeling during planarian regeneration. Nat Commun 14, 298. 10.1038/s41467-023-35937-y.

31. Sridhar, D., and Aboobaker, A. (2022). Monitoring Chromatin Regulation in Planarians Using Chromatin Immunoprecipitation Followed by Sequencing (ChIP-seq). Methods Mol Biol 2450, 529–547. 10.1007/978-1-0716-2172-1_28.

32. Dattani, A., Kao, D., Mihaylova, Y., Abnave, P., Hughes, S., Lai, A., Sahu, S., and Aboobaker, A.A. (2018). Epigenetic analyses of planarian stem cells demonstrate conservation of bivalent histone modifications in animal stem cells. Genome Res 28, 1543–1554. 10.1101/gr.239848.118.

33. Robb, S.M., and Sanchez Alvarado, A. (2014). Histone modifications and regeneration in the planarian Schmidtea mediterranea. Curr Top Dev Biol 108, 71–93. 10.1016/B978-0-12-391498-9.00004-8.

34. Jaber-Hijazi, F., Lo, P.J., Mihaylova, Y., Foster, J.M., Benner, J.S., Tejada Romero, B., Chen, C., Malla, S., Solana, J., Ruzov, A., and Aziz Aboobaker, A. (2013). Planarian MBD2/3 is required for adult stem cell pluripotency independently of DNA methylation. Dev Biol 384, 141–153. 10.1016/j.ydbio.2013.09.020.

35. Oviedo, N.J., Nicolas, C.L., Adams, D.S., and Levin, M. (2008). Establishing and maintaining a colony of planarians. CSH Protoc 2008, pdb prot5053. 10.1101/pdb.prot5053.

36. Shibata, N., and Agata, K. (2018). RNA Interference in Planarians: Feeding and Injection of Synthetic dsRNA. Methods Mol Biol 1774, 455–466. 10.1007/978-1-4939-7802-1_18.

37. King, R.S., and Newmark, P.A. (2018). Whole-Mount In Situ Hybridization of Planarians. Methods Mol Biol 1774, 379–392. 10.1007/978-1-4939-7802-1_12.

38. Stubenhaus, B., and Pellettieri, J. (2018). Detection of Apoptotic Cells in Planarians by Whole-Mount TUNEL. Methods Mol Biol 1774, 435–444. 10.1007/978-1-4939-7802-1_16.

39. Barghouth, P.G., Rojas, S., O’Dell, L.R., Betancourt, A.M., and Oviedo, N.J. (2022). Analysis of DNA Double-Stranded Breaks Using the Comet Assay in Planarians. Methods Mol Biol 2450, 479–491. 10.1007/978-1-0716-2172-1_25.

40. Oviedo, N.J., and Levin, M. (2007). smedinx-11 is a planarian stem cell gap junction gene required for regeneration and homeostasis. Development 134, 3121–3131. 10.1242/dev.006635.

41. Merighi, A., Gionchiglia, N., Granato, A., and Lossi, L. (2021). The Phosphorylated Form of the Histone H2AX (gammaH2AX) in the Brain from Embryonic Life to Old Age. Molecules 26. 10.3390/molecules26237198.

42. Peiris, T.H., Ramirez, D., Barghouth, P.G., Ofoha, U., Davidian, D., Weckerle, F., and Oviedo, N.J. (2016). Regional signals in the planarian body guide stem cell fate in the presence of genomic instability. Development 143, 1697–1709. 10.1242/dev.131318.

43. Podhorecka, M., Skladanowski, A., and Bozko, P. (2010). H2AX Phosphorylation: Its Role in DNA Damage Response and Cancer Therapy. J Nucleic Acids 2010. 10.4061/2010/920161.

44. Burma, S., Chen, B.P., Murphy, M., Kurimasa, A., and Chen, D.J. (2001). ATM phosphorylates histone H2AX in response to DNA double-strand breaks. J Biol Chem 276, 42462–42467. 10.1074/jbc.C100466200.

45. Lee, Y.J., Son, S.H., Lim, C.S., Kim, M.Y., Lee, S.W., Lee, S., Jeon, J., Ha, D.H., Jung, N.R., Han, S.Y., et al. (2020). MMTR/Dmap1 Sets the Stage for Early Lineage Commitment of Embryonic Stem Cells by Crosstalk with PcG Proteins. Cells 9, 1190. 10.3390/cells9051190.

46. Fagerberg, L., Hallstrom, B.M., Oksvold, P., Kampf, C., Djureinovic, D., Odeberg, J., Habuka, M., Tahmasebpoor, S., Danielsson, A., Edlund, K., et al. (2014). Analysis of the human tissue-specific expression by genome-wide integration of transcriptomics and antibody-based proteomics. Mol Cell Proteomics 13, 397–406. 10.1074/mcp.M113.035600.

47. Gegner, J., Gegner, T., Vogel, H., and Vilcinskas, A. (2020). Silencing of the DNA methyltransferase 1 associated protein 1 (DMAP1) gene in the invasive ladybird Harmonia axyridis implies a role of the DNA methyltransferase 1-DMAP1 complex in female fecundity. Insect Mol Biol 29, 148–159. 10.1111/imb.12616.

48. Flores, K.B., Wolschin, F., and Amdam, G.V. (2013). The role of methylation of DNA in environmental adaptation. Integr Comp Biol 53, 359–372. 10.1093/icb/ict019.

49. Pellettieri, J., Fitzgerald, P., Watanabe, S., Mancuso, J., Green, D.R., Sanchez Alvarado, A. (2010). Cell death adn tissue remodeling in planarian regeneration. Developmental Biology 338, 76–85. 10.1016/j.ydbio.2009.09.015.

50. Hubert, A., Henderson, J.M., Ross, K.G., Cowles, M.W., Torres, J., and Zayas, R.M. (2013). Epigenetic regulation of planarian stem cells by the SET1/MLL family of histone methyltransferases. Epigenetics 8, 79–91. 10.4161/epi.23211.

51. Neiro, J., Sridhar, D., Dattani, A., and Aboobaker, A. (2022). Identification of putative enhancer-like elements predicts regulatory networks active in planarian adult stem cells. Elife 11. 10.7554/eLife.79675.

52. Poulet, A., Kratikiewicz, A.J., Li, D., Van Wolfswinkel, J.C. (2023). Chromatin analysis of adult pluripotent stem cells reveals a unique stemness maintenance strategy. Science Advances 9. 10.1126/sciadv.adh4887.

53. Duncan, E.M., Chitsazan, A.D., Seidel, C.W., and Sanchez Alvarado, A. (2015). Set1 and MLL1/2 Target Distinct Sets of Functionally Different Genomic Loci In Vivo. Cell Rep 13, 2741–2755. 10.1016/j.celrep.2015.11.059.

